# Visual crowding in albinism: Evidence for a cortical sensory deficit with oculomotor influences

**DOI:** 10.64898/2026.03.16.712204

**Authors:** Vijay K Tailor-Hamblin, Maria Theodorou, Annegret Dahlmann-Noor, Tessa M Dekker, John A Greenwood

## Abstract

**Purpose:** Foveal vision in individuals with albinism is impaired not only by reduced visual acuity but also by elevated crowding – the disruption of object recognition in clutter. Because albinism is characterised by both retinal underdevelopment and nystagmus (uncontrolled eye movements), it is unclear whether crowding is elevated primarily from image motion due to eye movements or an additional sensory deficit. To disentangle these factors, we examined the spatial and featural selectivity of foveal crowding in albinism, comparing performance with controls and prior data from individuals with idiopathic infantile nystagmus syndrome (IINS), where nystagmus occurs without retinal underdevelopment.

**Methods:** Adults with albinism (n=8) and age-matched controls (n=8; 19–49 years) identified the orientation of foveal Landolt-C targets. In Experiment 1, targets were presented alone or flanked horizontally or vertically to assess spatial selectivity. In Experiment 2, flankers were of the same or opposite contrast polarity to assess featural selectivity. Stimulus size was adaptively scaled using QUEST to estimate gap-size thresholds.

**Results:** Crowding was substantially elevated in albinism, relative to both controls and IINS. Experiment 1 revealed stronger crowding for horizontally than vertically positioned flankers in albinism, mirroring the predominant direction of nystagmic eye movements. In Experiment 2, opposite-polarity flankers did not reduce crowding, indicating an absence of selectivity for target-flanker similarity.

**Conclusions:** Foveal crowding in albinism is markedly elevated, with a nystagmus-related spatial anisotropy and a lack of featural selectivity. These characteristics suggest that these elevations reflect both retinal image motion and a substantial sensory deficit arising from abnormal visual development.

## Introduction

Albinism is a genetic condition caused by reduced melanin production or dysfunctional melanin transport within the melanocyte containing cells ^1^, which arises within the first 6 months of life. Associated disruptions to oculo-motor development mean that a key clinical consequence of albinism is the presence of involuntary, uncontrolled eye movements. Albinism is accordingly classified in the over-arching infantile nystagmus syndrome ^2^, which also includes Idiopathic Infantile Nystagmus Syndrome (IINS), where the eye movements have no obvious physiological cause. Like IINS, the eye-movement instability in albinism is typically in the horizontal plane, with far less movement vertically, and with varied types of nystagmus waveforms including pendular or jerk movement patterns ^3^. Clear structural differences in retinal architecture nonetheless exist between individuals with IINS and albinism: individuals with IINS have seemingly normal foveal development, whereas individuals with albinism have varying grades of foveal hypoplasia ^4^. This retinal underdevelopment is likely a key driver of the marked impairments in visual acuity in albinism, relative to individuals with IINS ^5–8^. Here we examined the downstream consequences of albinism for visual crowding, a core aspect of cortical visual function.

In addition to deficits in visual acuity, albinism is associated with disruptions to numerous aspects of visual function. Both Vernier hyperacuity and contrast sensitivity have been found to be impaired in albinism, with the greatest deficits for vertically oriented gratings and lines ^8,9^, which would be most disrupted by the predominantly horizontal eye movements. These deficits are most pronounced in the fovea, with performance becoming equivalent to typical levels in peripheral vision. On this basis, Wilson, Mets, Nagy and Kressel ^9^ argued that albinism produces an arrested development of foveal vision, with lasting consequences for the organisation of the cortical mechanisms underlying spatial vision. This fits with broader evidence for delays in foveal and retinal development ^4,10^, as well as disruptions to retinotopic properties in visual cortical regions, particularly in albinotic individuals with achiasma, where the optic nerve fibres do not cross over to the opposite brain hemisphere ^11–15^.

A key aspect of cortical visual processing is *visual crowding*, the disruption to object recognition in cluttered scenes ^16^. Crowding effects are pronounced in peripheral vision ^17,18^, with far smaller effects in the typical fovea ^19,20^. Elevations in foveal crowding occur in a range of clinical conditions, including amblyopia ^21,22^ and IINS ^7,23,24^. In albinism, elevated foveal crowding was first reported in a small sample of individuals with albinism who showed disproportionately high thresholds when Landolt-C elements were flanked vs. in isolation, with greater elevations than those reported in IINS ^7^. Elevated crowding has also been demonstrated in children with both IINS and albinism ^25^. More recently, Moshkovitz et al ^26^ showed that the elevated crowding in albinism persists under both scotopic and photopic light levels, unlike in typical vision, a pattern argued to reflect the atypical development of cortical visual areas.

Elevations in foveal crowding have been more extensively studied in IINS, though evidence for their underlying mechanisms is mixed, with support for both a long-term sensory deficit in visual cortex and for more momentary effects related to eye movements, such as image smear or the displacement of elements into peripheral vision. Evidence for momentary effects comes from measurements of the spatial extent of foveal crowding in IINS, where individuals with IINS show greater elevations with horizontally than vertically placed flankers, matching the predominantly horizontal pattern of their eye movements ^24^. This anisotropy did not arise in typical adults and those with amblyopia, who showed equivalent crowding in the horizontal and vertical dimensions, but could be reproduced in typical participants with simulated nystagmus, adding further evidence for the momentary effect of eye movements. A similar pattern of spatial bias has been documented in other visual functions such as hyperacuity and contrast sensitivity in IINS ^27,28^, as is also the case in albinism ^8,9^.

Other aspects of crowding in IINS do nonetheless suggest the presence of a cortical sensory deficit. Firstly, several studies observe that the elevations in crowding induced by simulated nystagmus eye movements do not reach the same magnitude as that observed in people with IINS ^23,29^, suggesting at least some contribution from a sensory deficit. Second, we have recently found that the feature selectivity of crowding differs in IINS relative to typical peripheral vision. Crowding is typically strong when target-flanker similarity is high and reduced when similarity is low ^30,31^, particularly when elements differ in contrast polarity ^32–35^. On the contrary, individuals with IINS showed no improvement in crowded gap thresholds when target-flanker similarity was reduced via opposite contrast polarities ^29^. This pattern differed in simulated nystagmus, where performance matched that of typical peripheral vision. Altogether, the spatial pattern of crowding in IINS, which follows the trajectory of nystagmic eye movements, coupled with these divergences in the magnitude and selectivity of crowding in IINS, suggests that the elevations in foveal crowding reflect both a long-term sensory deficit and the momentary effect of eye movements. A similarly mixed pattern of evidence is also apparent for other visual deficits in IINS, including orientation discrimination ^36,37^, bisection ^38^, and vernier acuity ^39^.

To date, these properties of foveal crowding in albinism have not been examined, making the origin of the elevations unclear. Given the substantial evidence for both retinal and cortical disruptions in albinism, it is likely the elevations in crowding should be driven by long-term sensory deficits, at least in part. However, the presence of nystagmic eye movements in albinism, with a predominantly horizontal waveform like that found in IINS, are also likely to contribute. To disentangle these effects, we therefore examined the spatial and featural selectivity of crowding in albinism. Where prior studies have tended to group individuals with IINS and albinism together ^7^, here we sought to examine their visual abilities in isolation, under conditions that can be directly compared to those of IINS. We first examined the spatial profile of foveal crowding. Given the predominantly horizontal pattern of eye movements, we predicted that the spatial pattern of crowding in albinism would be similar to that of IINS ^24^, with greater crowding from horizontally-than vertically-placed flankers. We then examined the featural selectivity of crowding, predicting that albinism should show a similar invariance to target-flanker similarity as observed in IINS, unlike simulated nystagmus, which is wholly driven by the momentary effects of stimulus motion ^29^. In both cases, we predicted that crowding elevations should be of a much greater magnitude than that seen in IINS, given the clear evidence for cortical deficits in albinism. Together, these patterns would suggest a similar origin for crowding in albinism as that proposed for IINS.

## Experiment 1: The spatial selectivity of crowding

We first examined the spatial selectivity of crowding in individuals with albinism. Using Landolt-C elements, we compared performance with an isolated target (unflanked) and when two Landolt-C flankers were positioned horizontally or vertically around the target. In our prior work ^24^, individuals with nystagmus showed greater crowding when flankers were positioned horizontally compared to vertically, following the same pattern of the nystagmic eye movements. Similar patterns were found when nystagmus was simulated in typical vision using stimulus motion, suggesting an origin of the deficits in the eye movements. However, the magnitude of unflanked gap thresholds was greater in IINS compared with simulated nystagmus, suggesting some form of sensory deficit. In examining these conditions in albinism, our hypothesis was that crowding in albinism shares a common basis with that of IINS. If this were true, then crowding in albinism should be greater with horizontally-placed flankers, compared to vertically-placed flankers.

## Methods

### Participants

Sixteen adults underwent a full Orthoptic examination to ensure they met the inclusion and exclusion criteria into one of two groups: albinism (n = 8, mean age 30.1 years), and controls (n = 8, mean age 29.5 years). All participants were aged between 19-49 years, with no known neurological conditions. Control participants were included if they achieved a best-corrected visual acuity (BCVA) in each eye of 0.20 logMAR or better, with no strabismus or nystagmus present. Control participants were excluded if there were any signs of neurological disease, strabismus, nystagmus, or BCVA that was worse than 0.20 logMAR. For the albinism group, a horizontal, vertical, or torsional nystagmus waveform had to be present with a clinical diagnosis of albinism (ocular or oculocutaneous albinism), and a BCVA of 1.00 logMAR or better with both eyes open. Strabismus was allowed in the albinism group, though because testing was binocular, any unilateral amblyopia would not have caused elevated performance in either of the unflanked and flanked conditions, due to suppression of the amblyopic eye ^40^.

### Apparatus

Experiments were programmed using MATLAB (The MathWorks, Ltd., Cambridge, UK) on a Dell PC running Psychtoolbox ^41,42^ and the Eyelink toolbox ^43^. Stimuli were presented on an Eizo Flexscan EV2736W LCD monitor, with 2560 x 1440-pixel resolution, 60Hz refresh rate, and a physical panel size of 59.67 x 33.56 cm. Monitor calibration was undertaken using a Minolta photometer, with monitor luminance linearised in software to give a maximum luminance of 150 cd/m2. Participant responses to stimuli were indicated by a keypad with tactile markers on the keys to identify the response options. An Eyelink 1000 (SR research, Ottawa, Canada) recorded the position of the dominant eye at a sampling rate of 1000 Hz. Calibration of eye position was undertaken for control participants using the inbuilt Eyelink calibration. For participants with albinism, we first attempted to use the inbuilt calibration. Where this was not possible, a custom calibration procedure with 5 points of fixation (and post-hoc calibration of the data) was used, as described in Tailor, Theodorou, Dahlmann-Noor, Dekker and Greenwood ^24^, and based on similar procedures for the calibration of individuals with nystagmus ^44,45^. Viewing distance was varied between participants based on their BCVA in the orthoptic examination: 7 albinism individuals were tested at 78cms, 1 albinism individual was tested at 200cms, and all controls were tested at 300cms. This ensured both adequate resolution to measure acuity and sufficient range on screen to measure the extent of crowding. All participants were tested binocularly and wore their refractive correction where required, with no correction made for presbyopia. Participants had their head positioned on a chin rest with a forehead bar to minimise head movement. Data was analysed in MATLAB and SPSS.

Target and flanker stimuli were Landolt-C letters presented at the centre of the screen, either in isolation (unflanked) or flanked by two Landolt-C elements positioned either horizontally or vertically around the target. Elements were presented at 99.6% Weber contrast against a mid-grey background (Figure 1). Participants identified the position of the gap of the Landolt-C (four alternative forced choice, 4AFC), which was either 45°, 135°, 225°, or 315° on each trial. Oblique orientations were chosen to ensure equivalent identification levels of the target in all orientations (particularly across the two flanker configurations), as described in Tailor, Theodorou, Dahlmann-Noor, Dekker and Greenwood ^24^.

**Figure 1.**
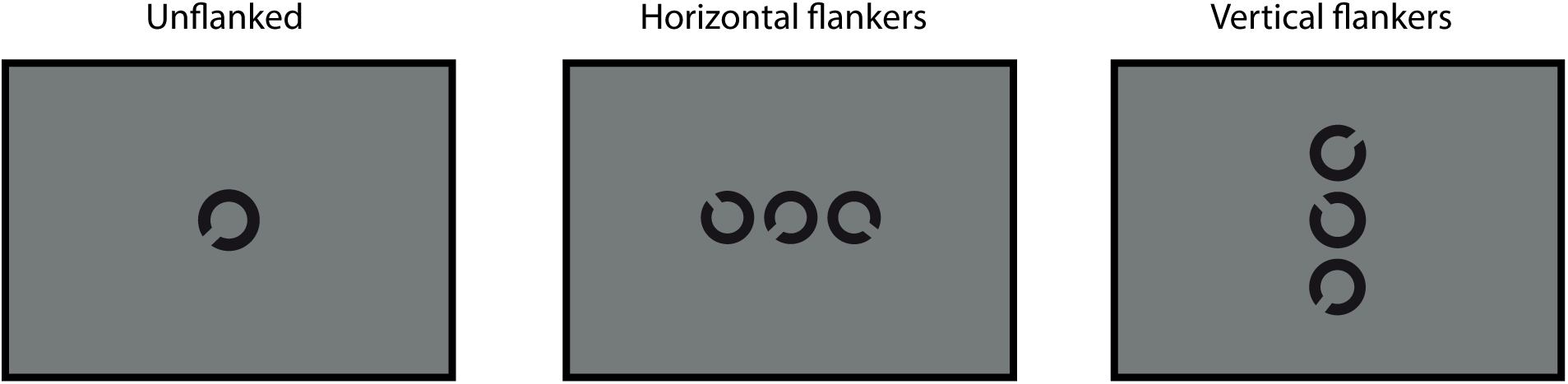
Example stimuli for the Landolt-C discrimination task in Experiment 1, using either unflanked (with an isolated Landolt-C), flanked-horizontal (where the target is the middle Landolt-C) and flanked-vertical arrangements.

Given evidence that nystagmats with IINS are “slow-to-see” ^46,47^, though cf. ^48^, stimuli were presented with unlimited duration until a response was made, at which point they were removed from the screen. Although this phenomenon has not been examined in individuals with albinism, the same procedure was retained to allow comparison across groups. Fixation guides were present throughout the experiment in the form of 4 white lines (the opposite polarity to the target) at a Weber contrast of 74.7%, located at the cardinal positions with a length of 50 pixels and with the inner edge separated from fixation by 15 times the stimulus diameter to avoid overlap. A 500ms inter-trial interval with a blank screen (leaving only a fixation point and the fixation guides) was then presented prior to the next stimulus. Feedback was not given. Participants were encouraged to make a choice in a timely manner.

The gap size and stroke width of the target and flanker Landolt-C elements was 1/5^th^ of the diameter, which was scaled using an adaptive QUEST procedure ^49^. The QUEST procedure converged on 62.5% correct (midway between chance and 100% correct). To avoid rapid convergence of the QUEST we added variance to the gap sizes presented on each trial by adding a value selected from a Gaussian distribution (with 0 mean and 1 SD) multiplied by 0.25 of the current trial estimate of the threshold. This minimised the number of trials presented at the same size in order to improve the subsequent fit of psychometric functions to the data ^50^. When flankers were present, their size matched the target, with a centre-to-centre separation from the target of 1.1 times their diameter, following the recommendations from Song, Levi and Pelli ^51^. With this scaling the effect of crowding can be measured using the elevation in threshold gap size in flanked conditions relative to unflanked thresholds. The constant scaling of both stimulus size and flanker spacing means allows the spatial extent of crowding to be quantified as the threshold gap size × 5 (stimulus diameter) × 1.1 (spacing).

At the start of each block of trials, participants began with 5 practice trials (identifying the orientation of the gap of the target Landolt-C), which were not included in the main analysis. Each block consisted of 60 trials (excluding practice trials), with 3 repeats per block to give a total of 180 trials for each stimulus condition (unflanked, horizontal flankers and vertical flankers). The whole experiment took approximately 1 hour. All procedures were approved by the NHS Northwest - Liverpool East Research Ethics Committee REC reference: 19/NW/0533. Participants were reimbursed for their time and travel expenses.

## Results and discussion

### Clinical demographics

Clinical characteristics can be found in Table 1. BCVA was measured using a logMAR chart, which gave a mean visual acuity (±1 standard deviation) of -0.03 ± 0.05 logMAR for the control group and 0.62 ± 0.17 logMAR for the albinism group. Control participants all demonstrated excellent stereo-acuity (measured with the Frisby stereo-test), with no ocular motility imbalances. None of the individuals with albinism demonstrated any 3D vision on the Frisby stereo test. All individuals with albinism showed nystagmus eye movements that were predominantly horizontal in direction, with 4 individuals showing a pendular waveform in the primary position. The others had a jerk waveform (n=2) or a periodic alternating nystagmus (n=2) where the waveform reverses its direction over time ^52^. Strabismus was present in 7 of the 8 individuals with albinism.

**Table 1.** Clinical characteristics of all participants. Key terms: RVA = right visual acuity, LVA = left visual acuity, BEO = both eyes open, NAD = no apparent deviation, LXT = left exotropia, RXT = right exotropia, LET = left esotropia, RET = right esotropia, F = full movement, BO = base out, BI = base in, CE = chin elevation, PAN = periodic alternating nystagmus. OCT grading was undertaken manually using the criteria from ^5^, N/A = not available

| Age<br>(years) | RVA<br>(logMAR) | LVA<br>(logMAR) | BEO<br>(logMAR) | Cover<br>Test | Ocular<br>motility | Head<br>Posture | Angle of<br>deviation | Stereopsis<br>(seconds<br>of arc) | Nystagmus<br>Type | OCT<br>grading |
| --- | --- | --- | --- | --- | --- | --- | --- | --- | --- | --- |
| **Control** |  |  |  |  |  |  |  |  |  |  |
| 38 | 0 | 0 | 0 | NAD | F | N | - | 55 | N | Normal |
| 39 | 0 | 0 | 0 | NAD | F | N | - | 55 | N | N/A |
| 28 | 0 | 0 | 0 | NAD | F | N | - | 55 | N | N/A |
| 27 | 0 | 0 | 0 | NAD | F | N | - | 55 | N | N/A |
| 19 | -0.1 | -0.1 | -0.1 | NAD | F | N | - | 55 | N | N/A |
| 23 | -0.1 | 0 | -0.1 | NAD | F | N | - | 55 | N | N/A |
| 24 | 0 | 0 | 0 | NAD | F | N | - | 55 | N | N/A |
| 38 | 0 | 0 | 0 | NAD | F | N | - | 55 | N | Normal |
| **Albinism** |  |  |  |  |  |  |  |  |  |  |
| 20 | 0.7 | 0.7 | 0.7 | RXT | F | N | 20BI | N | Jerk left | 3 |
| 26 | 0.6 | 0.7 | 0.6 | LET | F | N | 10BO | N | PAN | 4 |
| 39 | 0.74 | 0.64 | 0.64 | RET | F | N | 25BO | N | PAN | 4 |
| 33 | 0.7 | 0.85 | 0.7 | LET | F | N | 45BO | N | Pendular | 3 |
| 31 | 0.94 | 1 | 0.9 | LET | F | N | 30BO | N | Pendular | 4 |
| 38 | 0.6 | 0.8 | 0.6 | LET | F | CE | 25BO | N | Pendular | 3 |
| 27 | 0.78 | 0.98 | 0.7 | RET | F | N | 20BO | N | Pendular | N/A |
| 27 | 0.4 | 0.3 | 0.3 | NAD | F | N | 16BO | N | Jerk left | 4 |

Optical coherence tomography (OCT) was undertaken for the albinotic individuals to take cross-sectional images of the retina ^53^. The typical visual system shows a foveal pit and extrusion of the plexiform layers, with a lengthening of the outer segments of the photoreceptors and a widening of the outer nuclear layer (ONL) typically evident ^5^. Retinal grading of OCT images can be undertaken by examining the above 4 structures. Figure 2 shows example OCT images for 3 individuals: one with typical foveal and retinal structure, one with IINS (tested previously), where the presence of the structures in typical development are present, and one of the participants with albinism. In the albinism group, 3 participants had grade 3 foveal hypoplasia, with no foveal pit and a lack of widening in the cone OS. Four participants had grade 4 foveal hypoplasia (the highest level), characterised by an additional lack of widening of the ONL, as shown in Figure 2C. OCT scans were not available for the remaining participant. In the control group, typical foveal development was assumed, which was confirmed in 2 of the 8 controls with scans available (the remainder had no OCT scans available).

**Figure 2.**
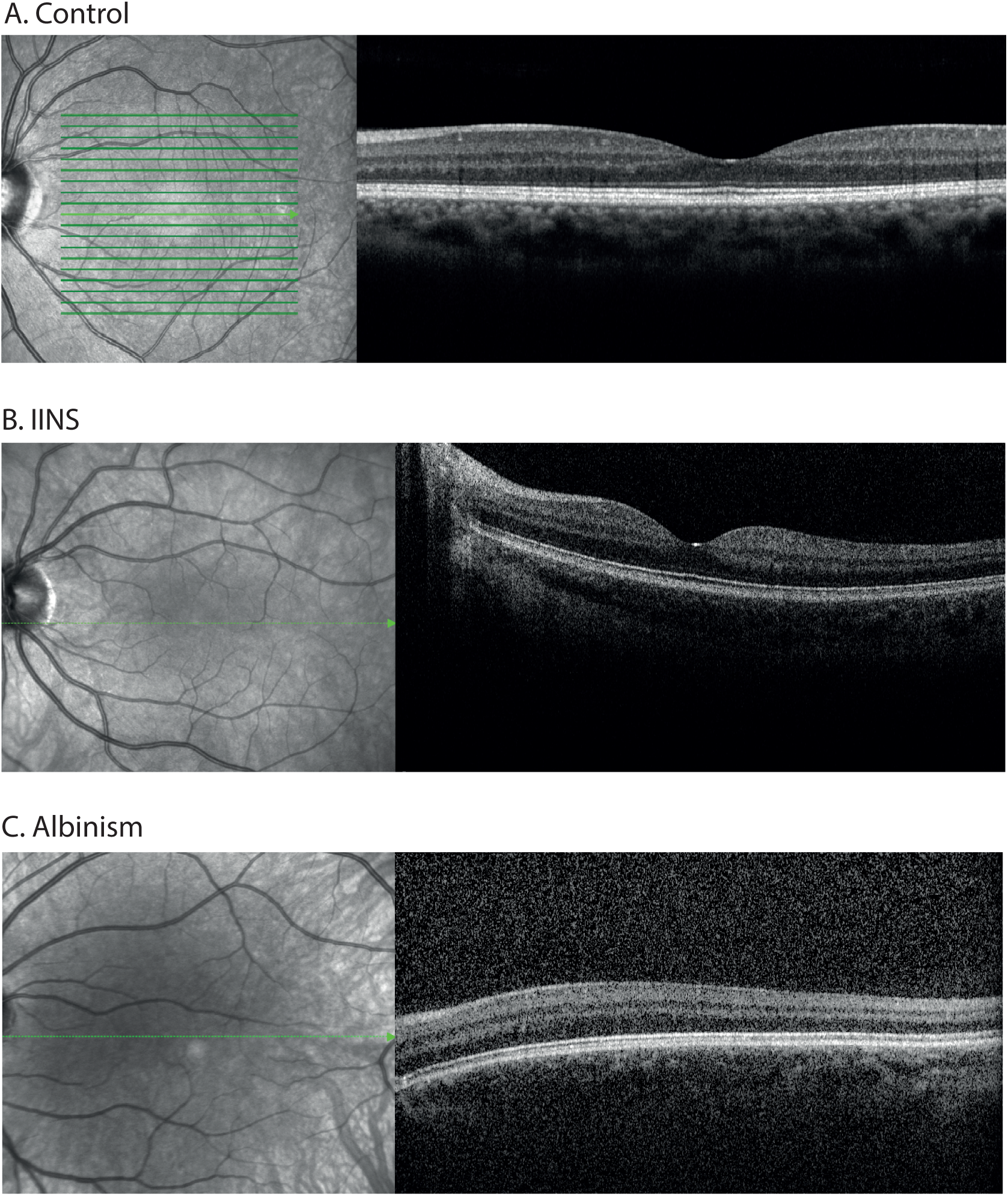
OCT images of the retina in 3 example participants. A. A typical adult, with normal foveal depression and all retinal layers present. B. An individual with IINS, classified as having normal foveal development. C. An individual with albinism showing grade 4 hypoplasia (no foveal architecture present).

### Eye movement characteristics

Recordings were made for all control participants, though due to difficulties in calibration and recording, fixational data was only obtained from 5 of the 8 individuals with albinism. To examine these eye movements, we first removed all blinks from recordings, plus 50ms before and after each blink. Saccades and micro-fixational eye movements were retained to characterise the full waveform of the eye movements during trials. To measure fixational variability, we first calculated the standard deviation (SD) of the horizontal and vertical position of the eye across all trials for each condition and took the mean of the SD across trials. These values were lowest in the controls for both the horizontal (0.43°) and vertical (0.47°) dimensions. Fixational instability increased for the albinism group, with a mean positional SD of 3.06° horizonally and 2.64° vertically. These values are higher than the variability of IINS individuals tested under the same conditions (horizontal = 1.60°; vertical = 1.08°) in both dimensions ^24^. Velocity values were calculated using a moving estimate of three samples to reduce error ^54^, before averaging across trials. Mean velocities were low in the control group, averaging 6.80°/sec horizontally and 8.89°/sec vertically. This rose substantially in the albinism group, with average velocites of 37.63°/sec horizontally and 19.33°/sec vertically. Altogether, fixational instability and eye-movement velocities were substantially elevated in the albinism group relative to controls, with more pronounced movements horizontally than vertically.

### Behavioural results

To analyse the behavioural data we combined repeated blocks of trials for each stimulus condition (unflanked, flanked-horizontal, and flanked-vertical) and participant, resulting in 180 trials per condition. For each stimulus size that was presented, proportion correct scores were collated. Psychometric functions were fitted to this data for each condition using a cumulative Gaussian function with 3 free parameters: midpoint, slope, and lapse rate ^55^. Because the QUEST gave variable trial numbers for each gap size, this fitting was performed by weighting the least-squared error value by the number of trials per point. Gap-size thresholds were derived from the psychometric function where performance reached 62.5% correct (mid-way between chance and ceiling) and converted to degrees of visual angle.

Figure 3A shows mean gap-size thresholds in minutes of arc for controls and participants with albinism, which were analysed using a 2×3 mixed-effects ANOVA with participant group (control, albinism) as a between-subjects factor and flanker condition (unflanked, flanked-horizontal, flanked-vertical) as a within-subjects factor. Analyses were run using logMAR thresholds (calculated using individual thresholds, to reduce heteroscedasticity in the data), as shown on the right side of the y-axis. The main effect of participant group was significant (F_(1,7)_ = 25.071, p < 0.001), with thresholds low for controls and clearly elevated for individuals with albinism across all conditions. The main effect of flanker condition was also significant (F_(2,14)_=88.420, p<0.001), though the interaction between flanker condition and participant group was not (F_(2,14)_=1.519, p=0.253). For controls, the unflanked condition gave the lowest thresholds, with modest elevations when both horizontal and vertical flankers were present, indicating a small crowding effect. Planned comparisons showed no difference between performance with horizontal and vertical flankers (t _(7)_ = 0.76, p=0.472). In the albinism group, unflanked thresholds were elevated relative to controls, with an additional elevation for both flanked conditions, indicating the presence of foveal crowding. Here, thresholds were significantly higher with horizontally placed flankers than vertically placed flankers (t _(7)_ = 3.01, p=0.020).

**Figure 3.**
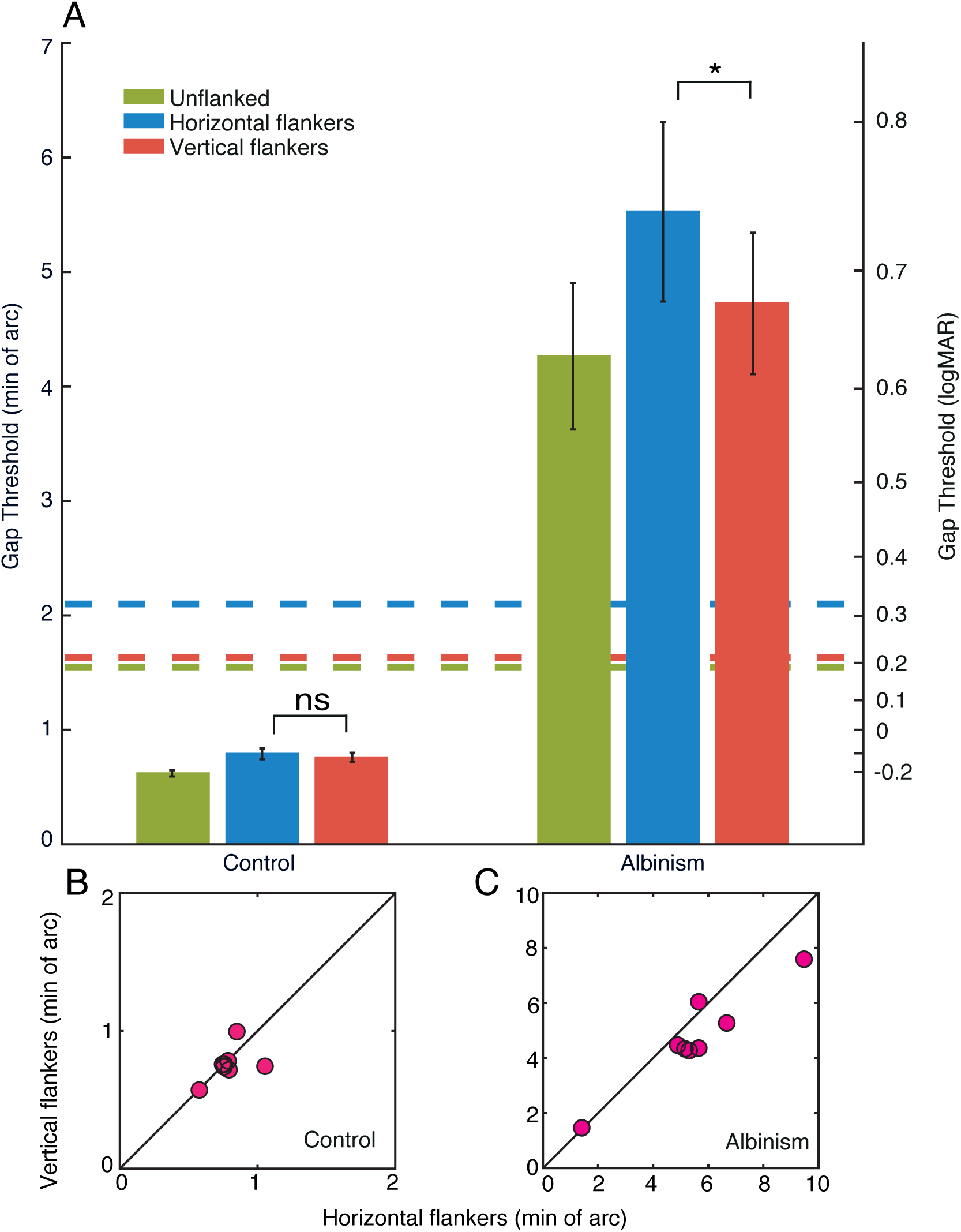
A. Gap-size thresholds for both groups, plotted in minutes of arc (left y-axis), with comparison to their logMAR equivalents (right y-axis). The blue bars plot unflanked thresholds, green bars the flanked-horizontal condition and red bars the flanked-vertical condition. Error bars represent the SEM, with *= significant and ns = not significant. Coloured dashed lines show average thresholds for the IINS group in Tailor, Theodorou, Dahlmann-Noor, Dekker and Greenwood ^24^. B. Gap thresholds (in min. of arc) for individual control participants, plotted for horizontal flankers on the x-axis against vertical flankers on the y-axis. The black line represents perfect correspondence between gap thresholds with horizontal and vertical flankers. Pink circles show individual participants. C. Gap thresholds for individuals with albinism, plotted as in panel B. Note the variation in X and Y scales.

Individual data in these flanked conditions further illustrate this difference between groups. For control participants (Figure 3B) thresholds with horizontal vs. vertical flankers clustered around the unity line, indicating little difference between these conditions. In contrast, 6 of 8 participants with albinism (Figure 3C) fell below the unity line, demonstrating consistently higher thresholds when flankers were horizontally than vertically placed.

The horizontal-vertical difference in the albinism group is similar to the pattern observed for individuals with IINS in Tailor, Theodorou, Dahlmann-Noor, Dekker and Greenwood ^24^, plotted in Figure 3A as dashed lines for comparison. Though both groups show the same anisotropy, with greater thresholds when flankers were horizontally vs. vertically placed, the albinism group showed larger thresholds in all conditions, indicating poorer overall performance. The level of performance in the albinism group was closer to that seen in the strabismic amblyopia group tested in Tailor, Theodorou, Dahlmann-Noor, Dekker and Greenwood ^24^, particularly in the unflanked condition, though crowding levels in albinism were slightly lower than in amblyopia. To quantify the elevation in these crowding levels relative to baseline acuity, multiples of crowding were calculated for each group by dividing flanked thresholds by unflanked thresholds. Expressed in this way, the elevation in thresholds was similar for the albinism and IINS groups. With horizontal flankers, thresholds were 1.30 times higher than unflanked acuity in albinism and 1.36 times higher in IINS. Vertical flankers gave smaller elevations of 1.11 times in albinism and 1.05 times in IINS. These values were smaller than the amblyopia group from Tailor, Theodorou, Dahlmann-Noor, Dekker and Greenwood ^24^, where thresholds rose 1.87 times with horizontal flankers and 1.62 times with vertical flankers. The level of crowding elevations relative to acuity loss is thus similar in albinism and IINS, while elevations are disproportionately large in amblyopia.

As outlined in the methods, the scaling approach used with our stimuli also allows for the estimation of the spatial extent of crowding ^51^. For controls, the spatial extent of crowding was small, with similar values of 0.072° with horizontal flankers and 0.069° with vertical flankers. These values were markedly increased in the albinism group, with an added anisotropy such that the horizontal extent was 0.507° and the vertical was 0.433°. As with the gap-size thresholds, these extent values exceed those reported in Tailor, Theodorou, Dahlmann-Noor, Dekker and Greenwood ^24^ for the IINS group (0.190° horizontally and 0.147° vertically) but remained slightly lower than those in amblyopia (0.634° and 0.549°).

In summary, our results show that individuals with albinism exhibit a pronounced horizontal elongation in the spatial extent of crowding, resembling the pattern observed previously in IINS and simulated nystagmus ^24^, and consistent with the predominantly horizontal nature of their eye movements. Threshold were however substantially higher in the albinism group for all conditions relative to both IINS and simulated nystagmus in Tailor, Theodorou, Dahlmann-Noor, Dekker and Greenwood ^24^, comparable to amblyopic levels, providing stronger evidence for an underlying sensory deficit in addition to oculomotor factors. At the same time, individuals with albinism showed higher fixational variability and speeds in their eye movements compared to those measured in IINS under matched conditions, which may contribute to some of the elevations in albinism. Overall, these findings indicate that crowding in albinism reflects a combination of image smear from unstable eye movements and a sensory deficit associated with structural abnormalities in the visual system.

## Experiment 2: The featural selectivity of crowding

In Experiment 1, the spatial selectivity of crowding in albinism indicated that both momentary eye-movement effects and a longer-term sensory deficit contribute to elevated crowding. Here we examined the featural selectivity of these effects. As outlined in the introduction, a key property of peripheral crowding is its selectivity for target-flanker similarity, where crowding is reduced when target and flankers are of opposite polarities ^30,32–34^. If the elevated crowding in albinism were primarily the result of image smear due to eye movements, these polarity-based modulations should not occur. Conversely, if the sensory deficit were to arise from retinal underdevelopment and associated reductions in foveal cell differentiation, the cortical regions processing foveal inputs may function more like those responding to peripheral vision. In this case, the selectivity of foveal crowding in albinism should match that of peripheral vision, with target-flanker differences in polarity producing improvements like those of peripheral crowding. A third possibility is that the sensory deficit in albinism differs from peripheral vision, producing elevated crowding without peripheral-like selectivity. Our recent examination of these effects in IINS was most consistent with this third mechanism: thresholds did not improve when the target and flankers had opposite polarities, unlike the advantage found in peripheral crowding, though the presence of target-flanker similarity effects in simulated nystagmus indicated that the momentary effects of eye movements produce distinct patterns ^29^.

We thus examined the effect of target-flanker similarity in albinism, comparing performance when all elements shared the same contrast polarity (all black or all white) or with opposite contrast polarities (target black and flankers white or vice versa). In line with a sensory deficit with properties distinct from crowding in peripheral vision, we predicted that target-flanker polarity differences will produce no improvement in thresholds for individuals with albinism. Note that we did not use temporal reversals of contrast polarity here, as in recent work ^29^, as these manipulations did not produce benefits for individuals with IINS.

## Methods

Participants for Experiment 2 were the same as in Experiment 1. The apparatus was also the same for both eye-tracking measures and the behavioural experiment. Target and flanker stimuli were again oriented Landolt-C letters, here presented either in isolation (target only) or surrounded by four flankers (positioned above, below, left, and right of the target) at the centre of the screen at 99.6% Weber contrast against a mid-grey background (Figure 4). As before, Landolt-C elements were scaled using a modified QUEST procedure, with proportions and inter-element spacing the same as in Experiment 1. Participants identified the location of the gap of the target Landolt-C, with four options (4AFC) of 45°, 135°, 225°, 315°, as before. On each trial, the target was either white or black (Figure 4A). For the two flanked conditions, the polarity of the target and flankers was either the same (flanked-same) as in Figure 4B, randomly all-black or all-white on each trial, or flanked-opposite, where the target and flankers had opposite polarity (Figure 4C), again with a random polarity for the target. Each of these three conditions was repeated three times for each participant.

**Figure 4.**
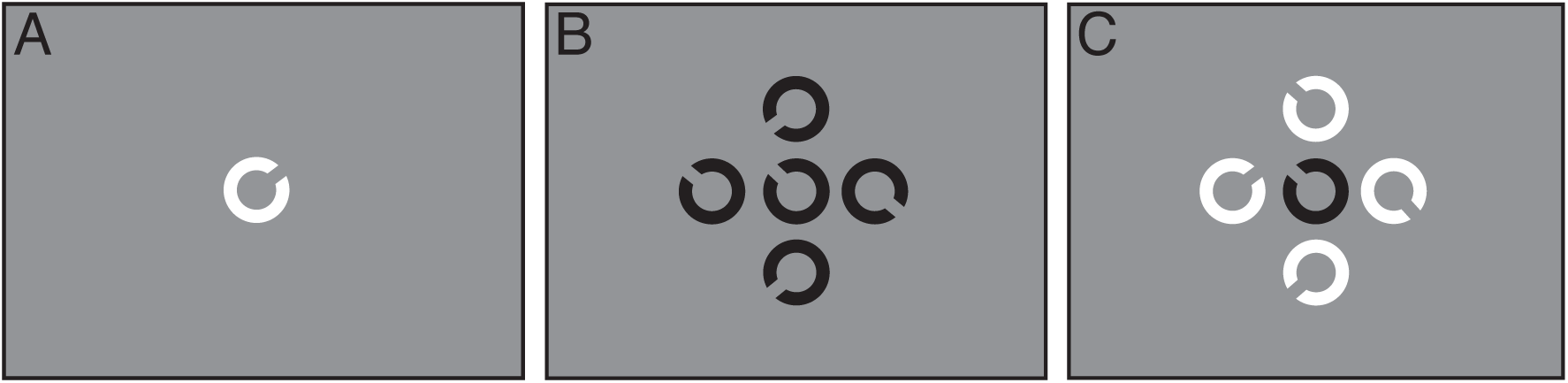
Example stimuli in Experiment 2. A. An unflanked target Landolt-C. B. The flanked-same condition, where a target Landolt-C (central element) was flanked with 4 elements sharing the same polarity. C. The flanked-opposite condition, where the target was flanked by 4 elements with the opposite polarity.

## Results and discussion

### Eye movement analysis

In this experiment, eye-tracking data was obtained from 6 of the 8 albinism participants, with difficulties in calibrating the remaining two. Given that participants were the same as in Experiment 1, analyses showed similar values for fixational instability and movement speed. For controls, the average variability in eye position was below 1° on both the horizontal (0.45°) and vertical (0.53°) dimensions. There was greater positional variability in the albinism group, with greater variability on the horizontal (3.26°) than the vertical (1.69°) dimension. This was again higher than individuals with IINS (1.31° horizontally and 1.05° vertically) under matched conditions ^29^. As in Experiment 1, average velocities were also low in the control group for both horizontal (7.09°/sec) and vertical (9.29°/sec) dimensions, which increased markedly in the albinism group, particularly on the horizontal dimension (32.32°/sec) relative to vertical (14.14°/sec). These values were comparable to our prior IINS group horizontally, where velocities averaged 30.46°/s, and were lower vertically where the average was 21.17°/s.

### Behavioural results

Analysis of the behavioural data was undertaken in the same way as in Experiment 1. Blocks of trials were combined for each participant and stimulus condition (unflanked, flanked-same, and flanked-opposite), resulting in 180 trials per stimulus condition. To examine these effects, a 2×3 mixed-effects ANOVA was conducted with participant group (control, albinism) as a between-subjects factor and flanker condition (unflanked, flanked-same polarity, flanked-opposite polarity) as a within-subjects factor, again using logMAR thresholds.

Figure 5A shows the mean gap-size thresholds for the control and albinism groups, plotted in minutes of arc (left y-axis) with corresponding logMAR equivalents (right y-axis). The main effect of participant group was highly significant (F _(1,7)_ = 78.889, p < 0.001), with larger thresholds in the albinism group than the controls. As in Experiment 1, thresholds in the albinism group were higher than those observed previously in individuals with IINS completing the same conditions (dashed lines), as reported in Tailor-Hamblin, Theodorou, Dahlmann-Noor, Dekker and Greenwood ^29^. The main effect of flanker condition was also significant (F _(2,14)_ = 49.433, p < 0.001), though there was no significant interaction between participant group and flanker condition (F _(2,14)_ = 2.396, p = 0.127). In the control group, gap thresholds were lowest in the unflanked condition, with a small but consistent elevation in thresholds for both flanked conditions, irrespective of polarity. Planned paired-samples t-tests confirmed that the difference between the two polarity conditions was not significant (t _(7)_ = 0.580, p = 0.580). Participants with albinism again showed higher levels of foveal crowding than the controls, with thresholds that were similarly high for both same- and opposite-polarity conditions and a non-significant difference between the two (t _(7)_ = −1.200, p = 0.269). The effect of target-flanker similarity was thus absent in both groups.

**Figure 5.**
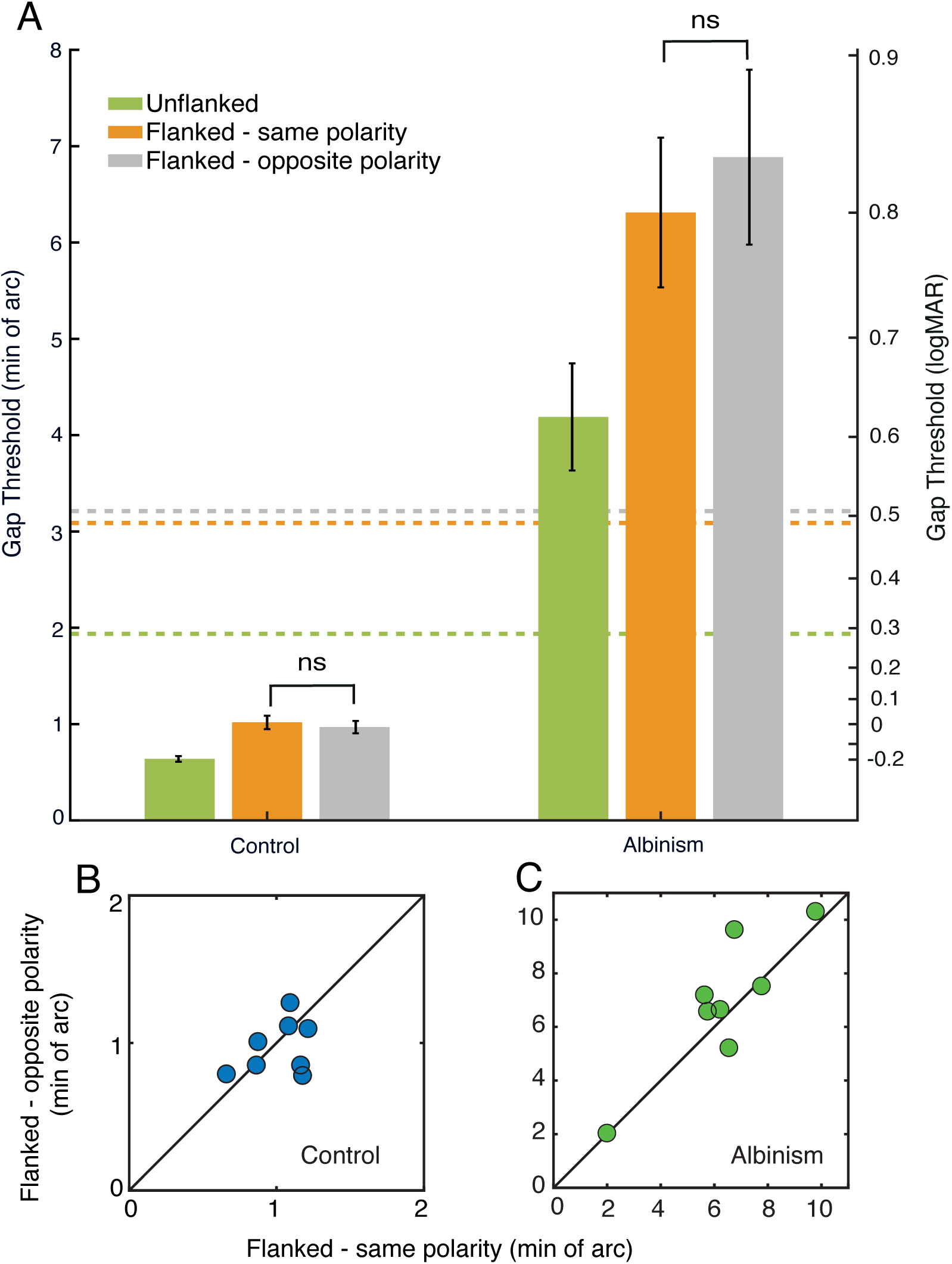
A. Gap thresholds for control and albinism participants in minutes of arc (left y-axis), with comparison to their logMAR equivalents (right y-axis). Green bars plot unflanked thresholds, orange bars the crowding-same condition and grey bars the crowding-different condition. Dashed coloured lines represent the corresponding thresholds for the IINS group from Tailor-Hamblin, Theodorou, Dahlmann-Noor, Dekker and Greenwood ^29^. Error bars represent the SEM, with *= significant and ns = not significant. B. Gap thresholds for flanked-same thresholds plotted on the x-axis, against flanked-opposite polarity thresholds on the y-axis for individual control participants. The black line shows perfect correspondence between conditions. Points show individual participants. C. Thresholds for individuals with albinism, plotted as in panel A. Note the variation in the X and Y scales.

To consider whether this null effect of target-flanker similarity was driven by outliers, we also examined individual thresholds, as plotted in Figure 5B for controls and 5C for the albinism group. In the control group, performance was evenly split, with 4 of 8 participants showing worse thresholds for same-polarity flankers and 4 out of 8 the opposite. In the albinism group, 5 of 8 participants showed worse thresholds in the flanked-opposite condition, and 3 the opposite. In neither case do we see evidence of any trend towards a release from crowding, further reinforcing the absence of a consistent polarity effect.

To summarise, gap-size thresholds were again elevated in participants with albinism relative to typical adults, both for flanked and unflanked conditions, as in Experiment 1. The magnitude of these elevations was again considerably larger than seen in individuals with IINS ^29^. In flanked conditions, the level of crowding was the same when flankers had the same contrast polarity as the target and when they differed, for both control and albinism groups. This is unlike crowding effects in typical peripheral vision, where opposite polarity produces improvements in performance ^30,32–34^, as well as in some prior studies examining foveal crowding effects ^32,56,57^, though not others ^58^; the opposite polarity condition for the albinism participants does not improve performance like that of the IINS individuals in Tailor-Hamblin, Theodorou, Dahlmann-Noor, Dekker and Greenwood ^29^. Altogether, the large elevations in crowding (beyond the levels of both IINS and simulated nystagmus), and the lack of selectivity for target-flanker similarity suggest the existence of a sensory deficit in albinism that operates in a manner that differs from the processes driving crowding in peripheral vision.

## General discussion

Our aim was to investigate the nature of the elevations in visual crowding in the fovea of individuals with albinism. Experiment 1 examined the spatial selectivity of these elevations, where participants with albinism showed a pronounced horizontal elongation of the interference zone for crowding, unlike in typical vision. This elongation had similar proportions to the elevations in foveal crowding seen in both IINS and simulated nystagmus ^24^, though the scale of interference was higher overall. Experiment 2 used variations in contrast polarity to examine the featural selectivity of these elevations. Unlike peripheral vision, where crowding is reduced by differences in the target and flanker elements, crowding in albinism remained elevated regardless of differences in contrast polarity. This pattern matches crowding in IINS, both of which differ from that of simulated nystagmus, where opposite contrast polarities reduced crowding ^29^. Altogether, our results suggest that elevations in foveal crowding arise through a combination of the momentary effects of eye movement and a longer-term sensory deficit.

Our observation that foveal crowding is elevated in albinism is consistent with prior findings ^7,25,26^. The larger magnitude of these elevations in albinism, relative to prior measurements in IINS ^24,29^, is also consistent with prior studies ^7,25^. It is possible that this difference may be partly driven by differences in the eye movement properties of the two groups, given the greater fixational instability and velocity we observe in albinism compared to IINS. However, the differences in these eye movement properties are smaller than the scale of the difference in thresholds, where performance in albinism is on average twice that of IINS. It is unlikely then that the momentary effect of these eye movements (such as image smear or peripheral relocation) could be the sole basis of the increased crowding for albinism, suggesting that an underlying sensory deficit must also drive these elevations.

A partial role for eye movements is also suggested by the spatial selectivity of crowding in albinism, which shows a horizontal elongation, consistent with the pattern in both IINS and simulated nystagmus ^24^. Similar performance anisotropies in albinism have been reported in a range of tasks ^8,9^, as in IINS ^27,28,36,38,39,59^. In the context of foveal crowding, elongations in the interference zone could relate to either the image smear of the target and flankers as the stimulus moves across the retina, or the relocation of the stimulus into peripheral retina where crowding increases. The results of Experiment 2 suggest that peripheral relocation is unlikely, since this should in theory give the selectivity for target-flanker similarity typically observed in the periphery ^30,32–34^, as observed in simulated nystagmus ^29^. It is possible then that image smear drives these effects, with the greater horizontal instability causing target and flanker signals to overlap, producing either masking effects by elements activating the same receptive fields over time ^60^. Alternatively, the long-term presence of these anisotropic movements may alter the shape of the pooling regions driving crowding, similar to suggestions regarding peripheral crowding ^61^. Regardless, the overall magnitude of both unflanked and flanked thresholds in albinism relative to IINS and the simulated effects of nystagmus in typical adults ^24^ suggests that eye movements alone are insufficient to produce the levels of foveal crowding in albinism.

The altered featural selectivity of foveal crowding in albinism adds further evidence that eye movements alone could not produce these deficits. In typical peripheral vision, crowding is reduced when target and flanker elements have opposite contrast polarities ^30,32–34^. In Experiment 2, we found no improvement in crowding when flankers had the opposite polarity to the target vs. when they were the same, matching the pattern observed in IINS ^29^. Complicating this picture is the similar lack of these target-flanker similarity effects in the typical fovea, both here and in our prior study using this same approach. Although this differs from some reports ^32,56,57^, others report a similar lack of selectivity for spatial frequency differences in foveal crowding ^58^. As above, this may be driven by eye movements activating the same receptive fields over time, causing masking effects whereby the flankers overwrite the target. The mixed effects of prior studies may reflect differences in stimulus arrangements, with some potentially less susceptible to target-flanker similarity effects than others (e.g. those that cover a larger expanse of the visual field such as arrays of vernier stimuli). Alternatively, the lack of effect in control participants may in part reflect limitations of the setup in the current study, where the low level of crowding in typical observers would make measurements of relative effects difficult. The higher level of crowding in albinism makes this unlikely to account for the null result in this group, however, particularly given the emergence of these effects in simulated nystagmus using the same apparatus ^29^. Regardless, the differences between foveal crowding in nystagmus and the typical periphery point to a sensory deficit in albinism that differs from that in other clinical conditions like amblyopia ^62^ where foveal crowding is similarly elevated. We suggest that the difference lies in the additional presence of nystagmic eye movements.

Several aspects of our findings point to foveal crowding in albinism being driven by a sensory deficit. As outlined in the introduction, albinism disrupts foveal development, often resulting in foveal hypoplasia ^4,10^. These disruptions have a downstream effect of cortical changes in both early and late visual cortex, with alternations in the pattern of retinotopy in visual cortical regions ^11–15^. It is likely that these disruptions are linked, given evidence for associations between retinal ganglion cell densities, cortical vision and crowding effects ^63^. A decline in neurons responding to the fovea, coupled with disarray in their retinotopic connections may increase the drive towards crowding effects in foveal vision. The presence of these cortical disruptions in albinism also explains the greater elevations than in IINS where the eye is structurally normal with minimal, if any foveal underdevelopment ^64^.

In drawing these conclusions, several limitations should be acknowledged. The sample size of our albinism cohort was modest, including participants with varying types and magnitude of nystagmus waveform, different degrees of foveal hypoplasia, and the presence of strabismus. Although our sample was sufficient to reveal clear group-level differences in foveal crowding, it is possible that some of these additional variables may have influenced the exact magnitude and pattern of selectivity seen in our results. Future work with larger, stratified samples could examine how crowding relates to the degree of foveal hypoplasia and to specific eye movement waveforms in particular. Difficulties in the calibration of the eye tracker also meant that eye-movement recordings were not available for all participants, limiting the strength of inferences between the properties of these eye movements and those of crowding. Our use of Landolt-C stimuli also provided a controlled way to quantify crowding, though it is possible that this does not fully capture the effect of visual disruptions to more complex, real-world visual tasks such as reading or object recognition. It nonetheless appears likely that crowding is a critical factor in shaping the everyday visual experiences of individuals with albinism. Finally, as the study was cross-sectional, we cannot determine whether the observed elevations in crowding reflect primarily congenital retinal underdevelopment, a long-term adaptation to nystagmus, or an interaction between the two.

In summary, our results demonstrated that crowding in albinism is characterised by three defining features: it is markedly larger in magnitude than in related conditions such as IINS, it exhibits a pronounced horizontal elongation that matches the anisotropy in eye-movement patterns, and it shows no selectivity for target-flanker similarity. The elevated magnitude of crowding in albinism relative to IINS in particular indicates that retinal image motion alone cannot account for these impairments, instead pointing to an additional sensory deficit, likely derived from retinal underdevelopment and associated downstream effects on visual cortical organisation. At the same time, the consistent horizontal elongation of the interference zone for crowding mirrors that observed in IINS and simulated nystagmus, implicating eye movements in shaping the spatial anisotropy of interference. Finally, crowding in albinism showed no benefit from target-flanker dissimilarity, unlike typical peripheral vision, suggesting that the sensory deficit is not simply peripheral-like but reflects a distinct alteration in central visual processing, similar to that observed in IINS. Together, these findings suggest a combined mechanism, whereby unstable eye movements produce the horizontal elongation and disrupt the effects of target-flanker similarity, while a longer-term sensory deficit governs the overall severity. With this dual contribution, retinal maldevelopment creates a predisposition for abnormal cortical organisation that is then further shaped by prolonged exposure to unstable visual input, leading to the pronounced and atypically tuned crowding observed in albinism.

